# Bayesian Multi-SNP Genetic Association Analysis: Control of FDR and Use of Summary Statistics

**DOI:** 10.1101/316471

**Authors:** Yeji Lee, Francesca Luca, Roger Pique-Regi, Xiaoquan Wen

## Abstract

Multi-SNP genetic association analysis has become increasingly important in analyzing data from genome-wide association studies (GWASs) and molecular quantitative trait loci (QTL) mapping studies. In this paper, we propose novel computational approaches to address two outstanding issues in Bayesian multi-SNP genetic association analysis: namely, the control of false positive discoveries of identified association signals and the maximization of the efficiency of statistical inference by utilizing summary statistics. Quantifying the strength and uncertainty of genetic association signals has been a long-standing theme in statistical genetics. However, there is a lack of formal statistical procedures that can rigorously control type I errors in multi-SNP analysis. We propose an intuitive hierarchical representation of genetic association signals based on Bayesian posterior probabilities, which subsequently enables rigorous control of false discovery rate (FDR) and construction of Bayesian credible sets. From the perspective of statistical data reduction, we examine the computational approaches of multi-SNP analysis using *z*-statistics from single-SNP association testing and conclude that they likely yield conservative results comparing to using individual-level data. Built on this result, we propose a set of sufficient summary statistics that can lead to identical results as individual-level data without sacrificing power. Our novel computational approaches are implemented in the software package, DAP-G (https://github.com/xqwen/dap), which applies to both GWASs and genome-wide molecular QTL mapping studies. It is highly computationally efficient and approximately 20 times faster than the state-of-the-art implementation of Bayesian multi-SNP analysis software. We demonstrate the proposed computational approaches using carefully constructed simulation studies and illustrate a complete workflow for multi-SNP analysis of *cis* expression quantitative trait loci using the whole blood data from the GTEx project.

## 1 Introduction

In the past decades, genetic association analysis has become a primary analytic tool to uncover genetic risk factors in complex diseases. With the advancement of high-throughput genotyping and phenotyping technology, genome-wide association studies (GWASs) and molecular quantitative trait loci (QTL) mapping have led to discoveries of an abundance of signals through genetic association analysis. These findings have subsequently played critical roles in exploring molecular mechanisms of complex diseases and predicting risks for individual patients.

Single-SNP association testing has long been considered as the standard approach for genetic association analysis. However, the results of the single-SNP analysis are not sufficiently informative by their own and often difficult to interpret without explicit references to linkage disequilibrium (LD) patterns of candidate variants. Additionally, it has been convincingly demonstrated that single-SNP testing fundamentally lacks power in identifying multiple association signals that are close by in relatively narrow genomic regions. A simple form of multi-SNP association analysis, known as *conditional analysis*, seeks a single “best” multi-SNP association model by a step-wise forward variable selection procedure (Yang et al., 2012). This approach addresses the power issue, but a single best solution oversimplifies the intrinsic difficulty introduced by the complex LD patterns and fails to account for the uncertainty of causal associations at SNP level.

Most recently, Bayesian approaches for multi-SNP association analysis have emerged as a promising alternative. They have at least two unique advantages over the traditional frequentist methods in the practice of genetic association analysis. First, they are built upon a natural hierarchical model that enables flexible incorporation of SNP-level functional annotations through principled prior specifications. Second, they utilize probabilistic quantification to characterize the strength of association evidence at SNP level, which can fully account for the complex LD structures presented in the genotype data. The successful applications of Bayesian genetic association analysis are illustrated in a wide range of applications for GWAS and molecular QTL mapping by piMASS (Guan and Stephens, 2011), GUESS (Bottolo et al., 2013), PAINTOR (Kichaev et al., 2014), CAVIAR (Hormozdiari et al., 2014), CARIVARBF (Chen et al., 2015) and FINEMAP (Benner et al., 2016), just to name a few. One of the significant limitations for the Bayesian approaches is the computational cost: instead of seeking a single best association model (i.e., by optimization), the Bayesian inference requires a comprehensive survey of all plausible association models (i.e., by integration). As a result, most existing Bayesian approaches do not scale well for extended genomic regions and often limited to the applications of fine-mapping analysis. Recently, we have proposed a new computational algorithm named deterministic approximation of posteriors (DAP), which is aimed to strike a balance between the commonly applied stochastic approximation algorithms (e.g., MCMC implemented in FINEMAP) and the exact computation by brute-force exhaustive search (e.g., in CAVIAR). We have shown, in Wen et al. (2016), that the DAP algorithm represents a highly efficient and accurate Bayesian inference procedure that can scale up to large-scale multi-SNP genetic association analyses in both GWAS and molecular QTL mapping.

Built upon the DAP algorithm’s high computational efficiency, this paper addresses two outstanding issues in the Bayesian multi-SNP genetic association analysis. First, we propose a novel false discovery rate (FDR) control procedure utilizing the posterior probabilities generated by our Bayesian approach. Rigorous control of type I error rate has always been an emphasis in genetic association analysis. Nevertheless, there is a lack of formal statistical procedures that can effectively control potential false discoveries in the multi-SNP analysis. Most theoretical results (Barber et al., 2015, Brzyski et al., 2017) on type I error control in the context of high-dimensional variable selection do not directly applicable to genetic association analysis because of the complex LD structures in the genetic association analysis. Our approach aims to fill this gap by proposing an intuitive hierarchical representation of association signals and adopting a principled Bayesian FDR control paradigm. Second, we discuss performing Bayesian multi-SNP association analysis based on summary statistics. Many authors have proposed association analysis algorithms that can work explicitly with summary-level data from single-SNP testing (Yang et al., 2012, Kichaev et al., 2014, Hormozdiari et al., 2014, Chen et al., 2015, Benner et al., 2016, Zhu et al., 2017). This has become an essential feature due to the nature of genetic data sharing for privacy protection. Our work on this topic focuses on understanding the analytic relationship of inference results based on individual-level data versus summary data. For example, we examine the following questions: do the two types of procedures (i.e., summary statistics vs. individual-level data) yield the same results? If not, is the inference based on summary statistics valid? Based on the answers to these questions, we attempt to identify a set of sufficient summary statistics that can lead to *identical* inference results as individual-level data, especially in Bayesian multi-SNP analysis.

The proposed novel computational approaches for multi-SNP genetic association analysis are implemented in the software package DAP-G, which is freely available at https://github.com/xqwen/dap/.

## 2 Method

### 2.1 Background, model and notation

In this paper, we focus on the problem of identifying potentially multiple genetic association signals using the following multiple linear regression model,

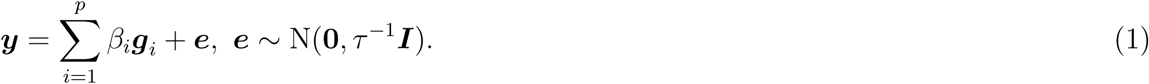

In practice, we assume that linear model (1) is obtained after regressing out a set of controlled covariates, including the intercept, from both the outcome vector and each genotype vector of candidate genetic variants. As a result, both ***y*** and all the ***g****i*’s have mean 0. Furthermore, we denote the *n × p* design matrix ***G***:= [***g***_1_ ***g***_2_ · · · ***g***_*p*_], which contains genotype data of all *p* candidate SNPs.

The point of interest for statistical inference is to identify the genetic variants that have non-zero effects on the quantitative trait. To this end, we explicitly define a latent binary indicator for each candidate predictor *i* by

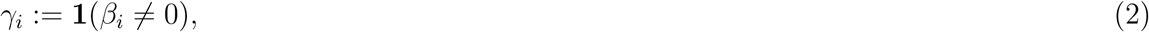

and 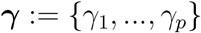

Based on this model, we formulate the problem of multi-SNP fine-mapping as a variable selection problem with respect to ***γ*** given the observed data (***y***, ***X***). Further details of the model are provided in Appendix A. Specifically, we compute the posterior probability for a given ***γ*** by

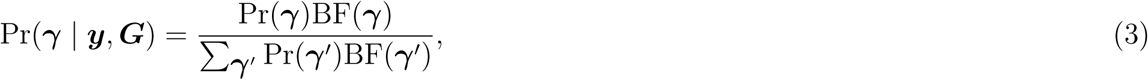

where Pr(***γ***) denotes the prior probability and 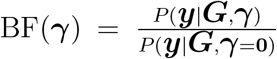 denotes the Bayes factor/marginal likelihood for ***γ***. Subsequently, the SNP-level posterior inclusion probability (PIP), which quantifies the strength of association for each SNP, can be marginalized from the posterior distribution, Pr(***γ*** *|* ***y***, ***G***).

#### 2.1.1 Overview of the DAP algorithm

For any given ***γ*** value, the prior and the Bayes factor can be analytically computed. The computational difficulty lies in the evaluation of the normalizing constant, i.e., 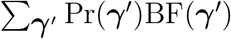: it is infeasible to enumerate all possible values of ***γ*** for a large number of candidate SNPs. The algorithm of deterministic approximation of posteriors (DAP) is designed to tackle this problem directly and can efficiently operate on a genomic region containing tens of thousands of candidate SNPs. (For larger regions or genome-wide analysis, it requires to segment the genome into LD blocks for separate processing.) The fundamental idea behind the DAP algorithm is based on the fact that noteworthy genetic association signals are typically *sparse* for any given genomic locus. Thus, only a very small number of candidate models (namely, the plausible models) make a substantial contribution to the normalizing constant. The DAP algorithm utilizes an efficient deterministic search strategy to identify the plausible models and approximates the normalizing constants based on the proven statistical principle known as *sure independence screening* (SIS,Fan and Lv (2008)). The approximation error to the true normalizing constant is also estimated in the search process, which plays a role in adjusting the estimated normalizing constant. In comparison, the commonly applied Markov Chain Monte Carlo (MCMC) algorithm is also designed to explore the plausible models but in a stochastic fashion. Because of the sampling space is enormous and consists of discrete models, it is unrealistic to expect that the MCMC algorithm reaches convergence with a limited computing resource. As a result, we find that the DAP algorithm often outperforms conventional MCMC algorithms in the setting of genetic association analysis. The new DAP-G algorithm is built upon the existing DAP algorithm and enjoys the same computational efficiency in the posterior inference of multi-SNP genetic association analysis.

### 2.2 False discovery rate control for genetic association signals

#### 2.2.1 Hierarchical representation of genetic association discoveries

Quantifying strength and uncertainty of genetic association signals is a long-standing problem in statistical genetics. The intrinsic difficulty lies in the fact that, with few exceptions, causal genetic associations may not be statistically identifiable at individual SNP level; Instead, each association signal is typically represented by a group of genetic variants whose genotypes are highly correlated. We argue that quantification and representation of a potential association signal should be dealt in a natural hierarchy, in which the following issues can be addressed:

1. the (un)certainty of the existence of an independent association signal;
2. the SNPs that are causally responsible for the association signal and their individual uncertainties

To demonstrate, we consider a hypothetical example from Wen et al. (2017): one of the two perfectly linked SNPs is causally associated with the complex trait of interest, and both SNPs are uncorrelated with the remaining candidate SNPs. In an ideal analysis, a precise characterization of the genetic association discovery should reflect that i) there is overwhelming evidence for the existence of an association signal; ii) there is a maximum degree of uncertainty to distinguish the causal variant between the two linked SNPs. We argue that the inference result of an ideal Bayesian analysis, which assigns PIP = 0.5 to each SNP, precisely encodes this information. The sum of the PIPs (= 1) indicates the sure existence of an association signal. Nevertheless, the two SNPs are equally likely to be the causal variant and not distinguishable solely based on the association data. (Further, if there exists additional information on the functional annotations of each SNP, it can be incorporated into the prior specifications that make the two SNPs distinguishable and potentially break the tie for the PIPs.)

This simple example illustrates the superiority of the probabilistic representation by Bayesian inference, which can carry comprehensive information from genetic association analysis. Nevertheless, we note that although almost all Bayesian multi-SNP analysis approaches generate SNP-level PIPs, there is no principled approach to summarizing the probabilistic evidence at the signal level, to the best of our knowledge. In a practical setting, it can be challenging to identify SNPs that are responsible for a single association signal (we will call the collection of such SNPs a signal cluster, henceforth). Identifying signal clusters require simultaneously examining both the overall evidence from multiple “similar” association models (e.g., when SNPs from the same signal cluster co-exist in an association model, the overall strength of evidence diminishes) and the pattern of LD.

The probabilistic quantification of association signals at both signal and SNP levels has multiple benefits. First, it allows rigorous control of the false discovery rate (FDR) at the signal cluster level (even though it can be challenging to pinpoint the causal association at the SNP level). Second, it allows constructions of Bayesian credible sets for suitable signal clusters, which is proven particularly attractive in genetic association analysis (Maller et al., 2012). Such credible sets provide a refined list of candidate SNPs for the underlying causal variants and can be critically valuable for the design of downstream molecular validation experiments.

#### 2.2.2 Identification of signal clusters

In the DAP-G algorithm, we integrate the functionality of automatic identification of signal clusters into the deterministic model search procedure.

Let ***γ***^(*i,j*)^, ***γ***^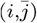^ and ***γ***^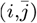^ denote three related association models that only differ in the values of *γ_i_* and *γ_j_*. Specifically, both SNP *i* and SNP *j* are assumed associated in ***γ***^(*i,j*)^, whereas only SNP *i* is assumed associated in ***γ***^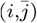^ and only SNP *j* is assumed associated in ***γ***^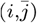^. We deem that SNP *i* and SNP *j* belong to the same signal cluster if and only if

1. the genotype *R*^2^ between SNP *i* and SNP *j* is greater than a pre-defined threshold;
2. the overall association evidence favors a single inclusion of SNP *i* or *j*, but not both, i.e.,

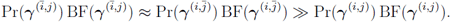

The first condition simply requires that SNPs within the same signal cluster are in LD and we use a rather relaxed threshold, i.e., *R*^2^ = 0.25, by default. In the second condition, 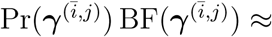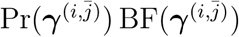 implies that the SNP *i* and SNP *j* makes similar contribution to the marginal likelihood with everything else being equal. However, when both SNPs within the same signal cluster co-exist in an association model, i.e., in ***γ***^(*i,j*)^, the likelihood is expected to be saturated, and the inequality is due to the prior “penalty” for assuming an additional causal SNP that is redundant. Essentially, this definition attempts to ensure that each signal cluster harbors precisely one independent association signal.

Based on above criteria, the DAP-G algorithm explicitly searches for redundant SNP representations of the same association signals and group them into signal clusters. When evaluating the approximate normalizing constant, each signal cluster is treated as an independent unit, and association models containing multiple SNPs from the same inferred signal cluster are explicitly avoided. We evaluate a signal-level PIP, denoted by SPIP, for each signal cluster by summing over the SNP-level PIPs from the member SNPs, i.e.,

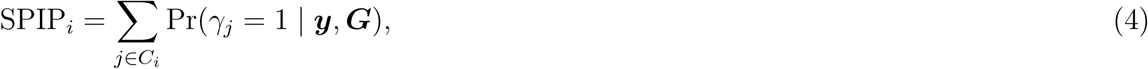

where the *C_i_* denotes the set of SNPs representing the *k*-th signal cluster. Note that our definition of the signal cluster and the search algorithm guarantees SPIP a valid probability distribution (i.e., strictly bounded by [0, 1]).

#### 2.2.3 Control signal-level false discovery rate

The signal-level PIPs enable a straightforward Bayesian FDR control procedure to guard against false positive findings. Specifically, the complement of SPIP is interpreted as the false discovery probability of signal cluster *i* and also known as the *local fdr* of the signal *i*, i.e.,

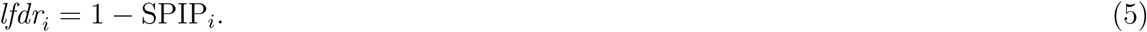

The use of local fdr for multiple hypothesis testing is well established in the statistical literature (Efron, 2012), and its result is asymptotically concordant to the frequentist testing approach utilizing *p*-values (Wen, 2018). Briefly, the following null hypothesis

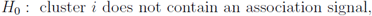

is rejected, if *lfdr_i_* is less than or equal to a pre-defined threshold *t*. Moreover, the threshold *t* is determined by the pre-defined FDR control level *α*, such that the average *lfdr* value from all rejected hypotheses is no greater than *α*. More precisely,

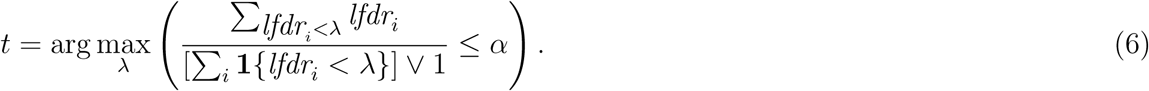

FDR control has become standard statistical approach for type I error control in molecular QTL mapping, where abundant association signals can be identified with modest sample size. Some also advocate direct specification of threshold value *t* (Efron, 2012), e.g., setting *t* = 0.05, which, in this case, is more stringent/conservative than controlling the overall FDR at 5%.

For a signal whose local fdr *≤ t*, it is straightforward to construct a (1 *− t*)% Bayesian credible set by selecting a minimum subset of SNPs, such that their cumulative SNP-level PIPs reaches 1 *− t*. The Bayesian credible intervals have been widely applied in GWAS since its introduction by Maller et al. (2012) in this context.

### 2.3 Inference using summary-level data

In many practical settings, individual-level genotype data may not be available, and association analyses have to rely on summary statistics. In this section, we discuss inference procedures to fit the proposed Bayesian hierarchical model utilizing only summary-level information. In comparison to the existing approaches in the literature (Chen et al., 2015, Zhu et al., 2017), we address this problem from a distinct point of view of statistical data reduction. In particular, we attempt to identify the *sufficient*, or near sufficient, summary statistics, which could potentially lead to a minimum or no loss of inference accuracy (comparing to using complete individual-level data). Moreover, we aim to examine if the commonly applied approaches, which utilize *z*-scores from single-SNP association testing, are optimal in multi-SNP association analysis.

The proposed Bayesian inference procedure depends on the observed genotype-phenotype data through the evaluation of the marginal likelihood, i.e., the Bayes factor,

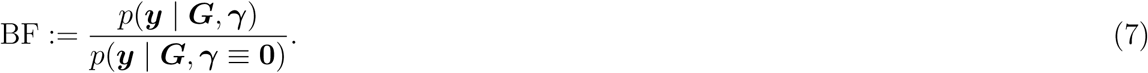

For an arbitrary ***γ***, Wen (2014) discusses an general analytic form of the Bayes factor with model (1) as a special case. (Note that, we take Wen (2014) as the starting point, because its results can be generalized to other designs of genetic association analysis, e.g., meta-analysis.) More specifically, if the residual error variance parameter *τ* is known, the analytic expression is exact; otherwise, it becomes an approximation by plugging in a point estimate of *τ*. The summary statistics required to compute the analytic form of BF include ***G****′****G*** (a *p × p* matrix), ***G****′****y*** (a *p*-vector) and a point estimate of *τ* if *τ* is not known (Appendix B.1). Under our formulation of the regression model (i.e., all ***g****i*’s are centered), the matrix ***G****′****G*** can be factored into

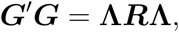

where ***R*** denotes the *p × p* sample correlation matrix between the *p* candidate SNPs, and **Λ** is a diagonal matrix defined by

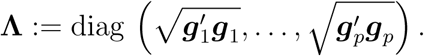

In the absence of individual-level data, some authors (Liu et al., 2014) have argued explicit sharing ***G****′****G*** for genomic regions of particular interests in multi-SNP fine-mapping analysis, many (Kichaev et al., 2014, Benner et al., 2016, Zhu et al., 2017) have proposed to estimate ***R*** and **Λ**, from an appropriate population panel. Henceforth, we assume that ***G****′****G*** is either provided or accurately estimated, and focus on the complete recovery of the information encoded in the *p*-vector, ***G****′****y***, from the summary statistics obtained in single-SNP testing.

In case that *τ* is known, we show that ***G****′****y*** can be accurately recovered given *z*-statistics and **Λ**.

This is because

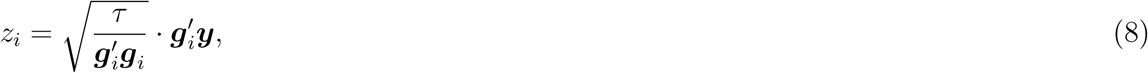

and

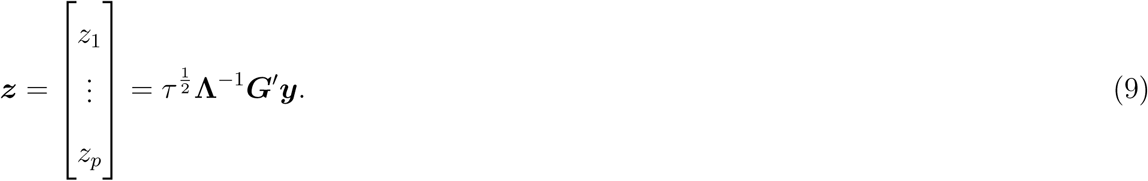

Therefore, it follow that

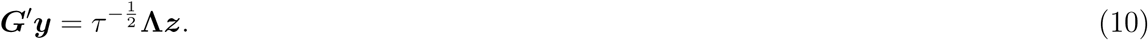

We note that Equation (9) directly leads to the *z*-score distribution utilized by FINEMAP and CaviarBF (Appendix B.3.1). For some specific type of normal priors on effect s *β*, which are explicitly scaled by Λ matrix, the required summary statistics can be reduced to (***R***, ***z***).

In practice, it is unrealistic to assume the knowledge of *τ* and *τ* is required to be estimated from the data. Note that even if the priors on genetic effects *β* are scaled by *τ*, as in the case of FINEMAP and CaviarBF, *τ* still explicitly enters into the Bayes factor computation (Appendix B.3.2). Precisely, Equation (9) should be modified to

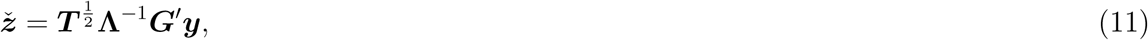

where ***T*** represents the following *p × p* diagonal matrix

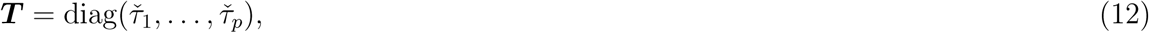

and each *τ̆_i_* represents the estimate of *τ* from the simple regression model testing the association of SNP *i*. For the first time, we provide rigorous justification to show that, under a specific prior specification, the summary statistics, (***R***, **Λ**, ***z****̆*), can be used to *approximate* required Bayes factor as an application of Laplace’s method (Appendix B.3.2). More importantly, our derivation and numerical experiments in Appendix B.3.2 also indicates that the residual error variance can be (sometimes severely) over-estimated in applying *z*-scores to approximate Bayes factors, especially when multiple independent signals co-exist. As a result, overestimation of the noise levels can lead to reduced power in uncovering true association signals.

To remedy the conservativeness of the *z*-score based inference procedure, we propose a new analytic strategy that enables more flexible and accurate approximation of marginal likelihood. Our approach requires following summary-level information,

1. estimated effect size and its standard error, (*b̂_i_*, se(*b̂_i_*)), from single-SNP analysis for each SNP *i* (note that, *z_i_* = *b̂_i_/*se(*b̂ _i_*));
2. sample size of the study, *n*;
3. total sum of squares (SST) of the quantitative trait: 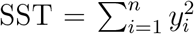, assuming ***y*** is pre-centered.

Let ***b****̂*:= (*b̂*_1_*,…,b̂ _p_*) and ***s****̂*:= (se(*b̂* 1)*,…*, se(*b̂_p_*)). We show that the complete summary statistics (***R***, ***b****̂*, ***s****̂, n*, SST) are sufficient to accurately recover ***G****′****y*** and ***G****′****G***. Furthermore, they allow estimating the corresponding MLE (or RMLE) of *τ* given ***γ***, which leads to a more accurate approximation of Bayes factors. The detailed justification and derivation are provided in Appendix B.2. The major benefits of the proposed approach are

1. It allows accurately estimating *τ* from the data matching any given ***γ*** value;
2. It allows work with an arbitrary type of the normal prior on genetic effect size (with or without scaling by *τ* and/or **Λ**).

Beyond the setting described by the model (1) for a single genetic association analysis, the proposed approach can be straightforwardly extended to multi-SNP analysis in a meta-analysis or trans-ethnic genetic association analysis using summary-level statistics Wen et al. (2015). (For this purpose, the second point above is particularly important.) From both simulations and real data analysis, we find that the ability to dynamically estimate *τ* according to the selected candidate SNPs can significantly improve the signal-to-noise ratios required for discovering multiple genuine genetic association signals. This factor likely explains the observation that approaches utilizing individual-level data typically outperform the existing approaches utilizing only *z*-scores. Our proposed strategy bridges this gap: if the LD information (namely, ***R***) is sufficiently accurate, the results based on the summary-level information are *identical* to those based on individual-level genotype data.

## 3 Results

### 3.1 Simulation studies

We set up a simulation scenario mimicking *cis*-eQTL mapping in a practical setting. In particular, we use the real genotype data from 343 European individuals from the GUEVADIS project (Lappalainen et al., 2013). We artificially construct a genomic region of 1,001 SNPs. The region is divided into 91 LD blocks, and each block contains 11 SNPs. All LD blocks are selected from chromosome 1, and the consecutive blocks are at least 1Mb apart. With this construction scheme, the LD only presents within each block, and the SNP genotypes are mostly uncorrelated across blocks (Supplementary Figure S1). We simulate a quantitative phenotype according to a sparse linear model. Specifically, with probability 0.05, an LD block is selected and a causal association is randomly assigned to one of its 11 member SNPs. On average, 4.75 genuine associations are expected from the whole region. The genetic effect of a causal SNP is independently drawn from a normal distribution, N(0, 0.6^2^), and the residual error for each sample is independently simulated from N(0, 1). Those particular parameters are selected such that the distribution of single SNP testing *z*-statistics from the simulated data matches the characteristics of the empirical distribution observed from the *cis*-eQTL analysis from multiple real eQTL data sets, namely GEUVDIS and GTEx (Supplementary Figure S2). We generate 1,000 independent data sets using this scheme.

The simulated data sets are analyzed using three methods:

1. DAP-G with sufficient summary statistics, i.e., (***R***, ***b****̂*, ***s****̂, n*, SST);
2. DAP-G with single SNP testing *z*-scores, i.e., (***R***, **Λ**, ***z****̆*);
3. FINEMAP with single SNP testing *z*-scores, i.e., (***R***, **Λ**, ***z****̆*)

The software package FINEMAP (Benner et al., 2016) implements a particular version of MCMC algorithm using the shotgun stochastic search scheme. Moreover, it utilizes the summary information (***R***, **Λ**, ***z****̆*) as input to compute the same approximate Bayes factors as in CAVIARBF. Because of its superior computational efficiency and accuracy compared to other available methods (see Benner et al. (2016) for details), we considered it the state-of-the-art for multi-SNP genetic association analysis using summary-level information.

We use the default priors for both DAP-G and FINEMAP, which are slightly different. DAP-G employs a more conservative default prior with respect to the simulated data sets, which assumes a single causal variant is expected *a priori*. FINEMAP, designed for fine-mapping analysis, assumes Pr(***γ*** = **0**) = 0 by default. In comparison, Pr(***γ*** = **0**) = (1 *−* 1/1001)^1001^ = 0.368 for DAP-G. As a result, we conclude that our simulated data scheme, in this case, slightly favors FINEMAP.

None of the methods assumes the knowledge of the artificial LD blocks constructed in the simulated data, i.e., LD information is inferred from the genotype data through ***R*** in all three approaches.

#### 3.1.1 Power for signal discovery

We first examine the power of all methods in uncovering the LD blocks that harbors a causal association signal.

Because the concept of a signal cluster is not defined in FINEMAP, we compute the cumulative PIPs for each constructed LD blocks and use this quantity to rank the blocks within each method. Although this approach does not always guarantee a valid probability for each pre-defined block, especially for FINEMAP, we find very few false positives from the blocks with cumulative PIPs *>* 1 for all three methods. We construct and compare the receiver operating characteristic (ROC) curves based on the ranking of the pre-defined LD blocks across all simulations. In addition to the three aforementioned approaches, we also rank the LD blocks by their the minimum *p*-values from the single SNP testing of their member SNPs. This approach is commonly used to identify eGenes (i.e., genes harboring at least an eQTL in their cis region) in *cis*-eQTL mapping.

Figure 1 shows the comparison of the ROC curves in a practically meaningful range, i.e., the false positive rate *<* 50%. In this set of simulations, the best performer is the DAP-G algorithm running with the sufficient summary statistics: for any false positive rate threshold, it always identifies more true positives than any of the approach in comparison. The difference in performance between the DAP-G algorithms using different summary statistics input confirms our theoretical argument for the superiority of the sufficient summary statistics over the *z*-scores. Although the DAP-G with *z*-score input and FINEMAP compute the approximate Bayes factor the same way, there is very noticeable difference reflected by the ROC curves. We suspect this difference is mainly attributed to the convergence issue of the MCMC algorithm employed in FINEMAP: if the MCMC run can be extended (significantly) longer, we expect these two sets of results would eventually converge. Finally, it is clear that all multi-SNP analysis approaches outperform the single-SNP method in identifying genetic loci that harbor association signals by a large margin in this simulation setting.

**Figure 1:**
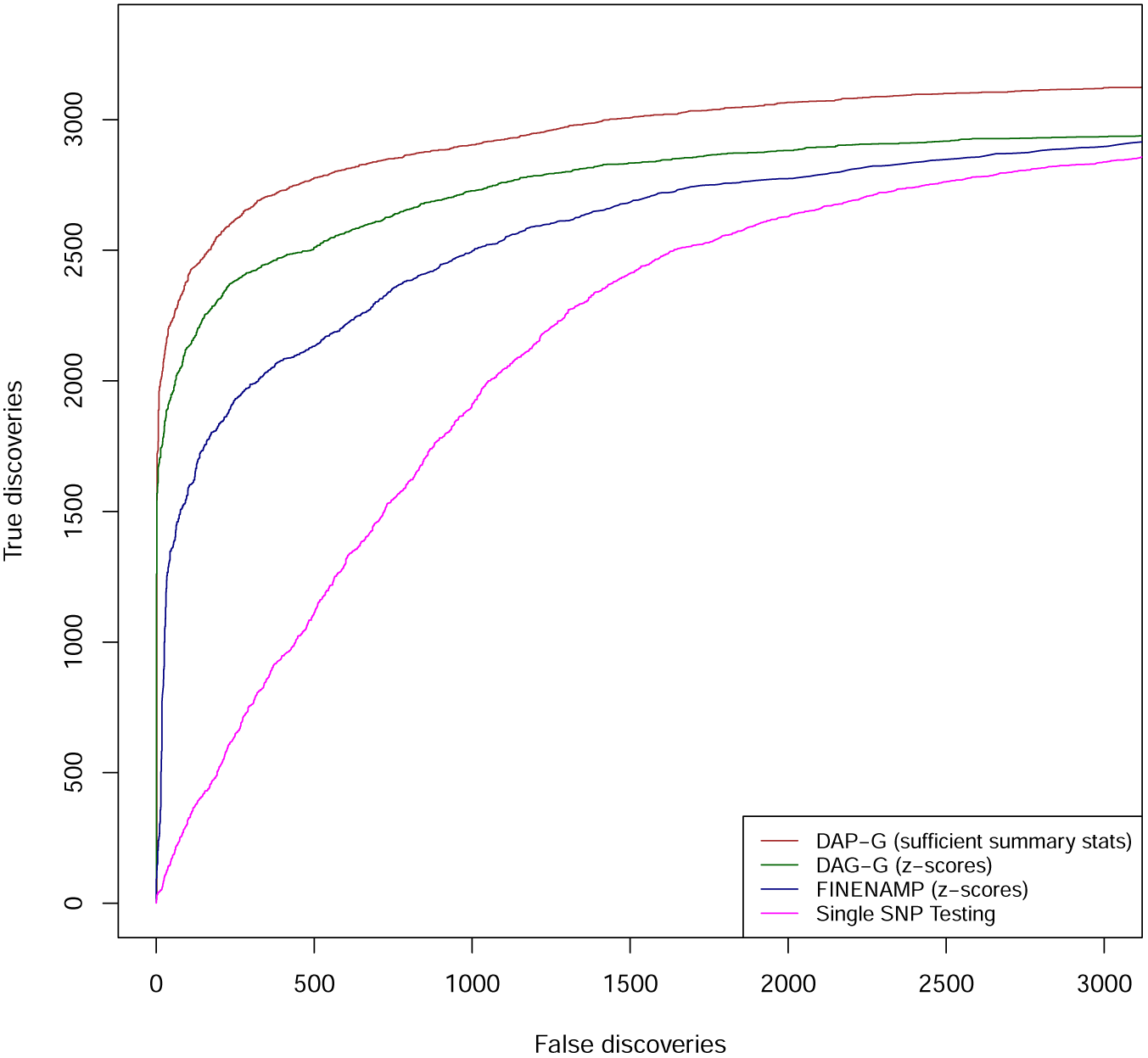
Power comparison in simulation studies. We examined the performance of 4 different methods in identifying the LD blocks that harbor true association signals. The methods compared include DAP-G using sufficient summary statistics (brown line), DAP-G using single SNP testing *z*-scores (dark green line), FINEMAP using single SNP testing *z*-score (navy blue line) and the single-SNP testing approach (magenta line). Each plotted point represents the number of true positive findings (of LD blocks) versus the false positives obtained by a given method at a specific threshold.

#### 3.1.2 Calibration of SNP-level PIP

Next, we inspect the calibration of SNP-level PIPs obtained from the different methods. The calibration of Bayesian posterior probabilities refers to the frequency property in repeated observations. For example in our specific context, it is expected that among many SNPs assigned PIP = 0.50, half of them are genuinely associated if the PIPs are indeed calibrated. The calibration of the posterior probabilities indicates the robustness of the model and the accuracy of the Bayesian computation.

For each method examined, we group all SNPs across simulated data sets into 10 bins according to their reported PIP values (namely, [0, 0.1), [0.1, 0.2)*,…*, [0.9, 1.0]). We then compute the proportion of truly associated SNPs in each bin. We expect that the frequency value is aligned to the average PIP value for each bin for calibrated SNP-level posterior probabilities.

Figure 2 shows that, among three methods compared, DAP-G running with sufficient summary statistics yield most calibrated SNP-level posterior probabilities. As expected, the PIPs by DAP-G using *z*-scores as input are slightly conservative. For FINEMAP, the posterior probabilities in some high-value PIP bins are shown to be anti-conservative, indicating potential convergence issues in MCMC runs.

**Figure 2:**
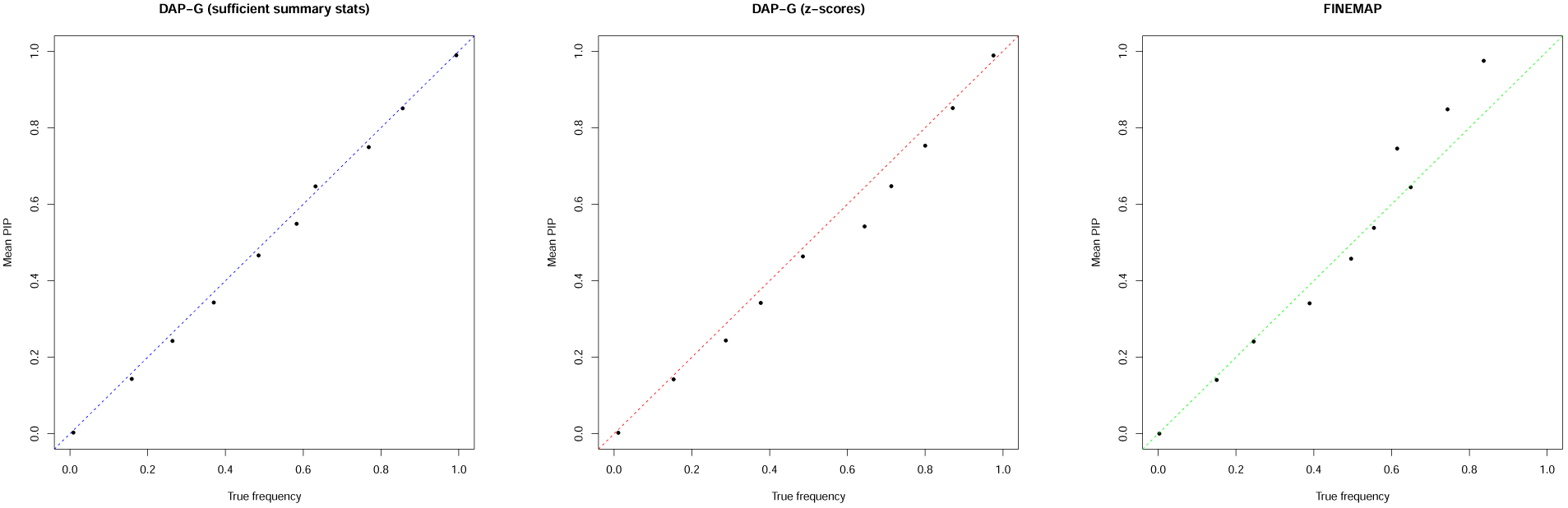
Calibration of SNP PIPs in the simulation study. PIPs from three Bayesian multi-SNP analysis methods (DAP-G with sufficient summary statistics, DAP-G with *z*-scores and FINEMAP with *z*-scores) are examined. PIPs from each method are classified into 10 equal-length frequency bins, the average PIP versus the corresponding true proportion (i.e., frequency) of causal SNPs for each bin is then plotted for each bin. If the PIPs are calibrated, we expect all points are aligned in the diagonal line. Points deviating from the diagonal line indicate that the PIPs may not be calibrated. More specifically, points below the diagonal line imply that the corresponding PIPs are conservative and points above the diagonal line suggest the PIPs are anti-conservative.

#### 3.1.3 FDR control at signal level

We then proceed to inspect the performance of FDR control at the signal cluster level by DAP-G. We performed the proposed Bayesian FDR control procedure using the inferred SPIP values. We label a true discovery if an inferred signal cluster indeed contains a causal SNP and the corresponding SPIP is greater than a pre-defined threshold and a false discovery otherwise. Subsequently, we compute the realized false discovery proportion (FDP) and the power with respect to the corresponding FDR control threshold. We repeat this procedure for a set of FDR levels ranging from 0.01 to 0.25. The detailed results are shown in Table 1. In all cases, the FDR’s of the signal clusters are conservatively controlled at all pre-defined levels. Furthermore, by utilizing sufficient summary statistics the power of discovering association signals is consistently higher than using *z*-scores.

**Table 1:** Realized signal-level false discovery proportion (FDP) and power in simula-tion studies. In all cases, the actual FDP values are below the target FDR control levels. As expected, the powers of DAP-G using sufficient summary statistics are consistently higher than the using the *z*-score based summary statistics.

| FDR control level | DAP-G (z-scores) |  | DAP-G (sufficient summary stats) |  |
| --- | --- | --- | --- | --- |
|  | FDP | power | FDP | power |
| 0.01 | 0.009 | 0.452 | 0.002 | 0.529 |
| 0.05 | 0.013 | 0.501 | 0.010 | 0.576 |
| 0.10 | 0.021 | 0.535 | 0.030 | 0.605 |
| 0.15 | 0.040 | 0.560 | 0.064 | 0.623 |
| 0.20 | 0.067 | 0.580 | 0.107 | 0.634 |
| 0.25 | 0.100 | 0.598 | 0.148 | 0.646 |

#### 3.1.4 Computational efficiency

Our implementation of the DAP-G algorithm is highly efficient: we observe that DAP-G runs magnitude faster than the state-of-the-art FINEMAP program. The speed-up is mainly due to the nature of deterministic search algorithm. Additionally, the implemented functionality of parallel processing for the DAP-G deterministic search procedure (via the OpenMP library) also contributes to the improved computational efficiency. For a dataset contains 5 independent signals, DAP-G runs about 1.5 seconds with 4 parallel threads and correctly identifies 4 of the 5 signals. In comparison, FINEMAP also achieves the same accuracy, and the runtime is benchmarked at 1 minute and 45 seconds on the same computer. The total user time for analyzing the complete set of 1,000 simulated data sets are 34 minutes 48 seconds and 741 minutes 6 seconds for DAP-G and FINEMAP, respectively. With 4 data sets being simultaneously analyzed on an eight-core Xeon 2.13 GHz Linux system, the real time of the complete analysis for DAP-G and FINEMAP are 7 minutes 50 seconds and 190 minutes 36 seconds, respectively.

### 3.2 Multi-SNP analysis of *cis*-eQTLs in GTEx whole blood samples

In this section, we illustrate a complete process of *cis*-eQTL mapping of the GTEx whole blood samples (version 6p) using the proposed DAP-G algorithm. The GTEx whole blood data include 338 individuals for which dense genotyping are performed. The expressions of 22,749 protein-coding and lincRNA genes are measured by RNA-seq experiments. The individual-level genotype-phenotype data are available for analysis. We followed the procedures described in GTEx Consortium (2017) to perform pre-processing and quality control of the genotype and expression data. For *cis*-eQTL mapping, we focus on the candidate genetic variants located within a 1Mb radius of the transcription start site (TSS) of each gene. On average, there are 7,118 candidate genetic variants per gene and no further SNP filtering procedure is taken before the multi-SNP association analysis.

We take an empirical Bayes approach to estimate the prior inclusion probability for each SNP. The estimation procedure, implemented in the software package TORUSWen (2016), utilizes the single-SNP association testing results across all genes. Additionally, it can incorporate SNP-level annotation data. We classify the candidate SNPs into 21 categories according to their distances to the TSS (DTSS) of corresponding genes and allow the priors vary in different categories. This decision is motivated by the previous observations (in almost all eQTL studies) that the abundance of *cis*-eQTLs is strongly associated with SNP DTSS. Our estimated priors by DTSS bins (Figure 3) from the GTEx data clearly confirms this pattern.

**Figure 3:**
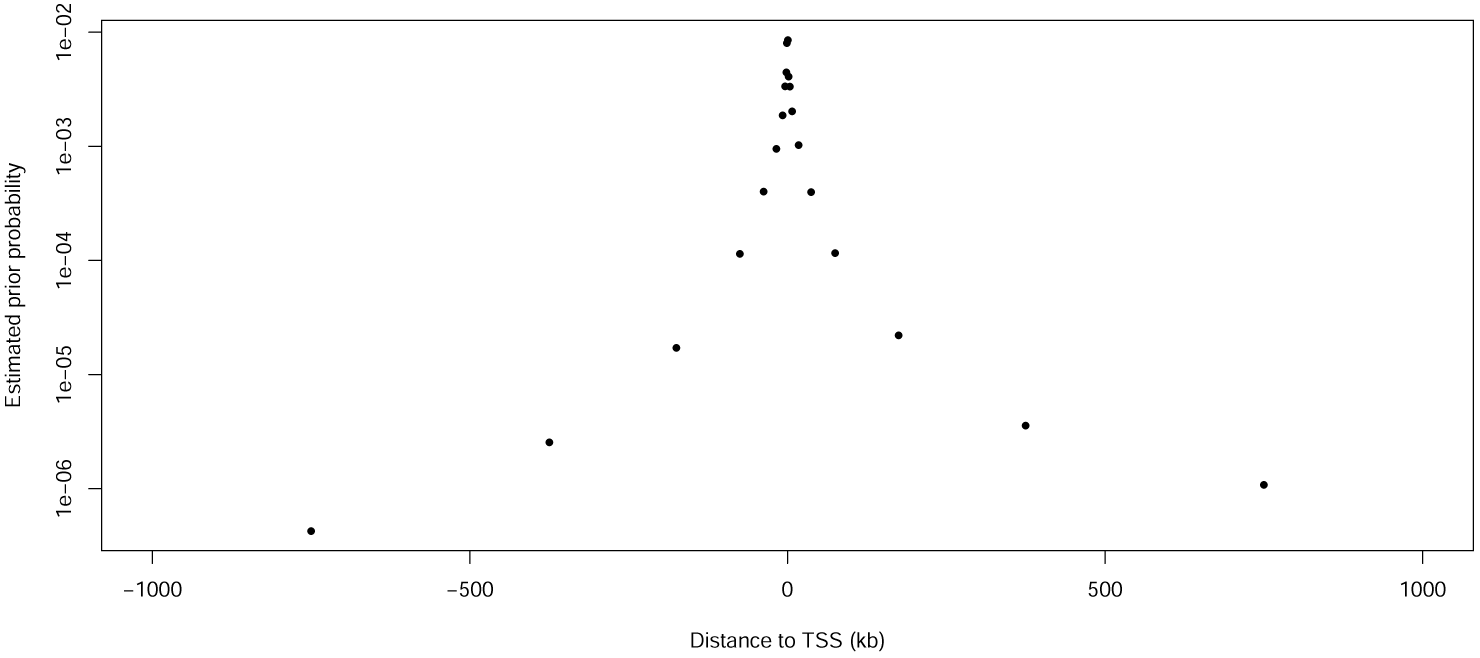
Relationship between estimated *cis*-eQTL priors and the SNP distances to transcription start sites (DTSS). All *cis* candidate SNPs are classified into 21 unequal-length bins according to their DTSS values. An EM algorithm implemented in the software package TORUS is used to estimate the prior inclusion probability for SNPs in each bin. Note that the quantitative distance information for the distance bins is *not* used by the EM algorithm. Each point on the plot represents the middle point of a distance bin, and its corresponding estimated prior. The result displays a clear pattern of fast decay of the abundance of eQTLs away from transcription start sites.

We then proceed to analyze all 22,749 genes using DAP-G. On a computing cluster and with 30 to 50 genes simultaneously analyzed, the processing of the complete data set takes about 14 hours. First, we compute the posterior expected number of *cis*-eQTLs for each gene by

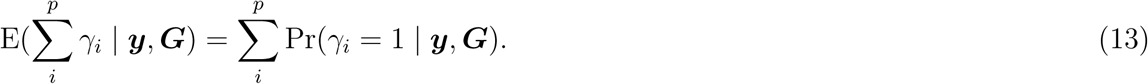

Figure 4 shows the histogram of the expected number of *cis*-eQTLs across all genes, which indicates that we are able to confidently identify multiple independent *cis*-eQTLs for a good proportion of genes. Applying the proposed FDR control procedure, we identify 9,056 independent *cis*-eQTL signals from 7,135 unique genes by controlling FDR at 5% level. A subset of 6,328 signals from 5,123 unique genes exceeds the more stringent threshold at 5% local fdr, for which we can construct 95% credible sets. There is a substantial variation in the size of the 95% credible sets (Figure 5). The median size of the credible sets is 7, and the mean is 14.9. The average pairwise *r*^2^ between SNPs in a credible set is 0.85 (median = 0.89). The largest credible set observed in this data set represents a *cis*-eQTL signal for gene *KANSL1* (ensembl id: ENSG00000120071) located at chromosome 17 (SPIP *~* 1.0), which consists of 354 tightly linked SNPs (average pairwise *r*^2^ = 0.90). Even for a single gene, we sometimes observe various sizes of credible sets. Figure 6 shows gene *TMTC1* (ensembl id: ENSG0000133687) for which we confidently identify 4 independent *cis*-eQTL signals. Interestingly, two of the signals have relatively small 95% credible sets containing 1 and 4 SNPs, respectively; while the credible sets for the other two signals are noticeably larger, containing 20 and 32 SNPs, respectively. These results reinforce our observations that causal associations can be complicated to identify even if the evidence for the existence of an association signal (e.g., SPIP) is overwhelming.

**Figure 4:**
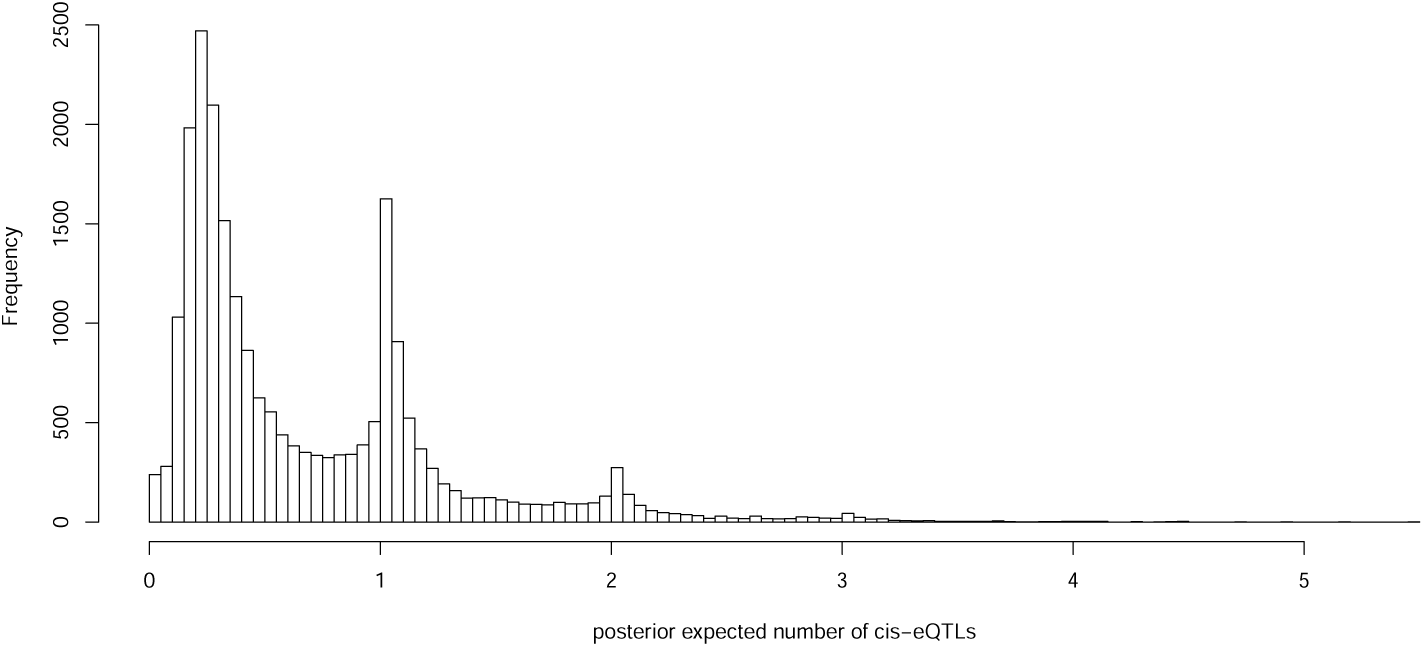
Histogram of posterior expected number of *cis*-eQTLs for 22,749 protein-coding and lincRNA genes analyzed in the GTEx whole blood data.

**Figure 5:**
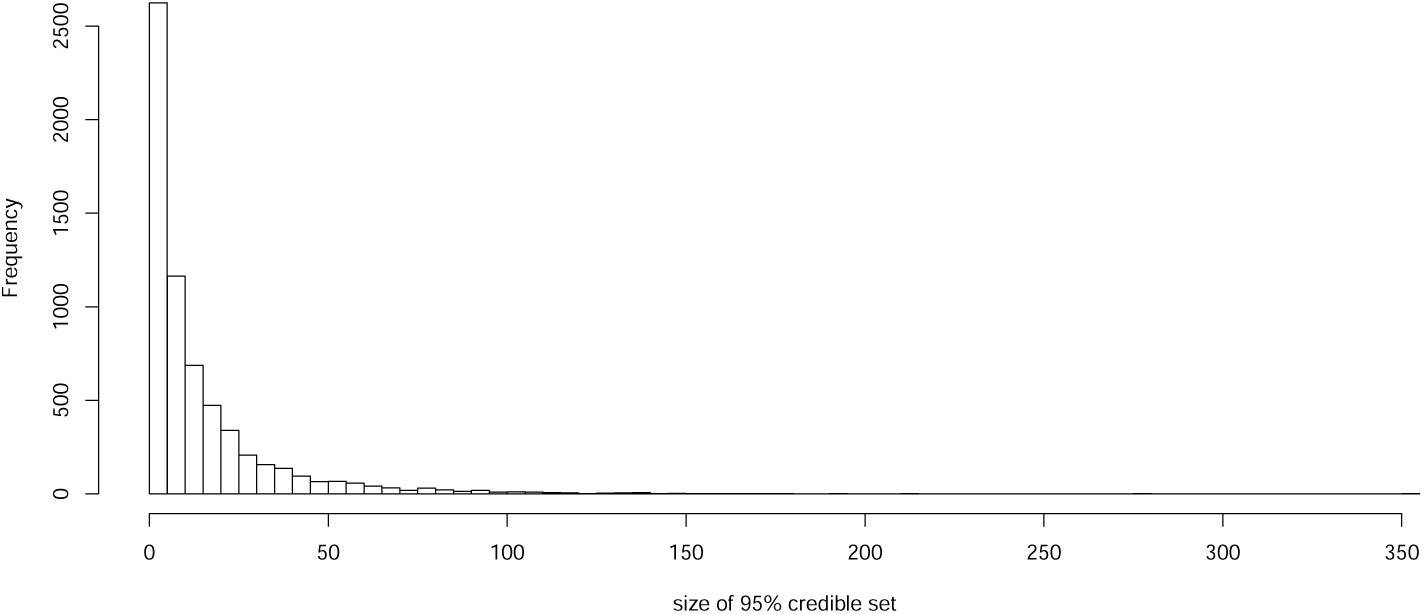
Histogram of the size of 95% credible sets constructed for 6,328 independent whole blood *cis*-eQTLs using GTEx samples.

**Figure 6:**
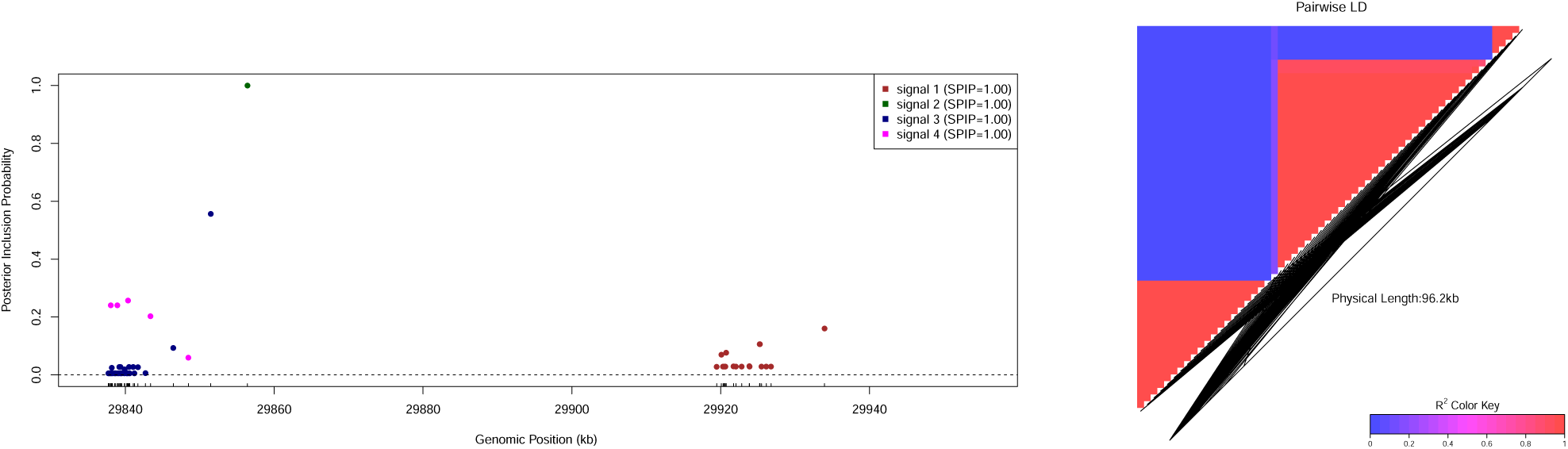
*cis*-eQTLs identified for gene *TMTC1*. The left panel shows 4 independent association signals are confidently identified in the *cis*-region of gene *TMTC1* (all SPIPS *→* 1). Each colored point represents a member SNP in the corresponding 95% credible set. The size of the credible sets differs according to different LD patterns. The right panel plots the LD pattern (*R*^2^) between the plotted SNPs. There is high LD within each signal cluster and very weak LD between the clusters.

To compare results with summary statistics based inference, we extract summary-level information from the complete data in two forms: the sufficient summary statistics, (***R***, ***b****̂*, ***s****̂, n*, SST), and commonly used (***R***, ***z****̂*). As predicted by our theoretical arguments, we find that the inference results based on sufficient summary statistics are identical to the analysis of individual-level data, whereas noticeable discrepancy can be observed from the inference results applying *z*-score based summary statistics. Figure 7 shows the comparison of SNP-level PIPs among the three different inputs for gene *TMTC1*. Particularly when *z*-scores are used as input, we note that the SPIPs for the third and the fourth signals in the original analysis for *TMTC1* are severely under-estimated and the 95% credible sets can no longer be constructed. In comparison, the SPIPs for the first two signals are still close to 1, and the corresponding credible sets mostly remain the same. We find these results are also consistent with our observations from the simulation studies.

**Figure 7:**
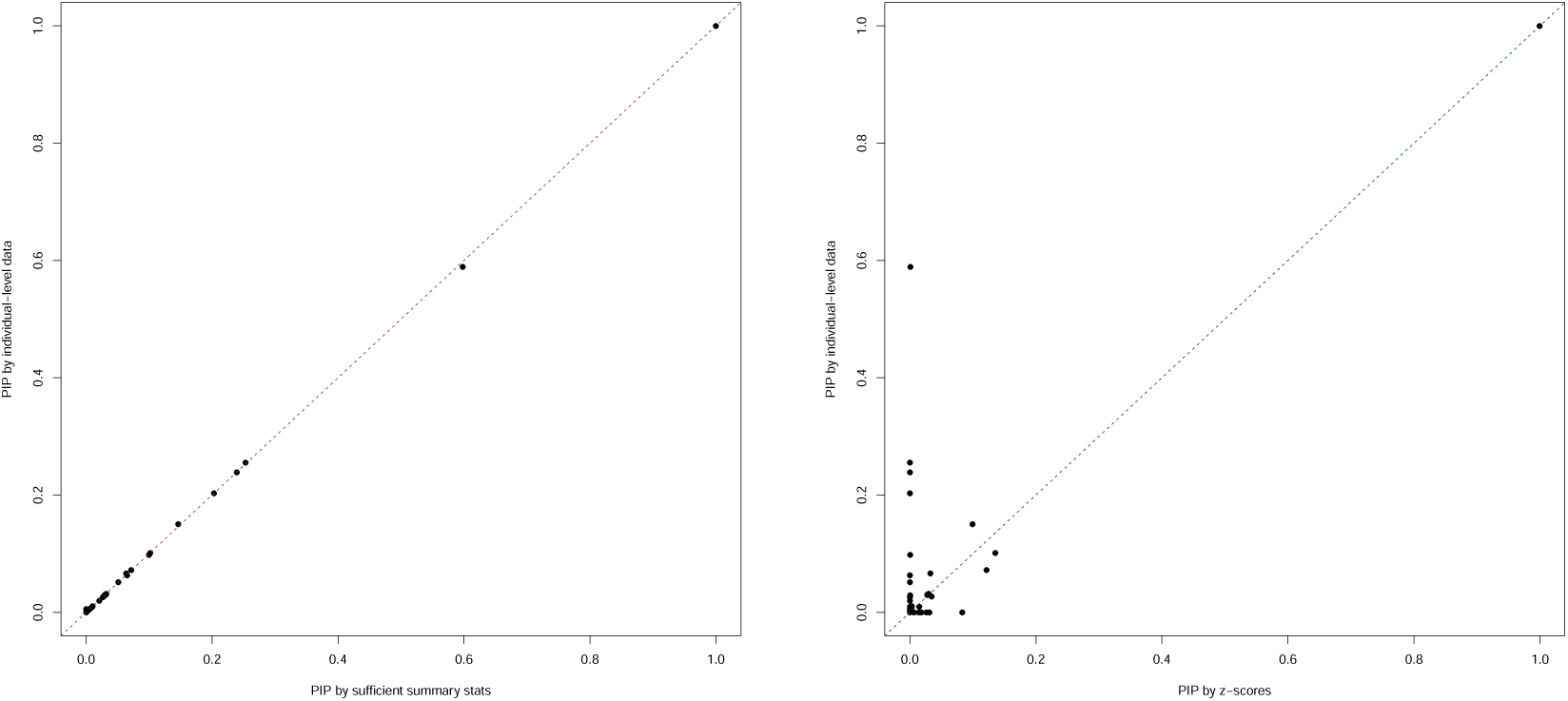
Comparison of PIPs computed from individual-level data versus summary statistics. The PIPs for 863 *cis* candidate SNPs for gene *TMTC1* are plotted. All PIPs are computed by DAP-G. The left panel shows the PIPs computed from sufficient summary statistics, and they are identical to the PIPs computed from individual-level data. The right panel shows the PIPs computed from *z*-scores, which are noticeably conservative, for most cases, in comparison to the PIPs computed from the individual-level data.

In summary, we find our *cis*-eQTL mapping analysis by DAP-G is highly efficient. The multi-SNP analysis results are more informative and more natural to interpret in comparison to the standard single-SNP analysis. We provide the complete analysis results in, which include the quantification of all*cis*-eQTL signal clusters and corresponding credible sets.

## 4 Discussion

In this paper, we have described a powerful and efficient computational approach to perform multi-SNP genetic association analysis. Within the Bayesian framework, we have introduced a new paradigm to comprehensively represent a complex genetic association signal in a natural hierarchy that accounts for LD structures and easy to interpret. With the probabilistic quantification of the strength of association evidence, we have shown rigorous FDR control can be straightforwardly applied. From the perspective of data reduction, we have derived the sufficient summary statistics, (***R***, ***b****̂*, ***s****̂, n*, SST), that result in identical inference with individual-level data in quantitative trait mapping. Furthermore, we are also able to establish the theoretical connection to the commonly applies inference based on summary-level data, which is shown to be a conservative approximation to the exact inference using individual-level data.

In *cis*-eQTL mapping, we have illustrated that multi-SNP analysis can completely replace the need for reporting single SNP analysis findings because of its informativeness and efficiency. We believe the same argument can be made regarding the analysis of GWAS data. We acknowledge that almost all multi-SNP genetic association analysis approaches, including ours, do not computationally scale beyond a genomic region up to 4 Mb and most commonly applied for fine-mapping analysis instead of genome-wide scan. Many have shown (Berisa and Pickrell, 2016, Wen et al., 2016) that it is effective to apply a divide-and-conquer strategy that segments genome according to population-specific LD blocks and performs multi-SNP analysis independently on each LD block. This strategy may be necessary if genomic annotations are incorporated into GWAS analysis and an unbiased enrichment analysis contrasting annotated functional SNP versus unannotated is desired. Additionally, with sample sizes of GWAS reach to the bio-bank scale, improved power for uncovering modest genetic association signals has become critically important. As shown in our simulation study, especially Figure 1, identifying critical regions through filtering via single SNP testing may not be the best practice.

With the previous results on computing Bayes factors in complex linear systems (Wen, 2014), our results presented in this paper can be straightforwardly extended to accommodate many different study designs for studying complex and molecular traits. The important applications include multi-SNP analysis in meta-analysis setting and eQTL mapping across multiple tissues, just to name a few. With the availability of the analytic forms of approximate Bayes factors under these complicated settings, it is now possible to perform rigorous FDR control and carry out the computation through summary statistics.

Genetic association analysis is not and should never be the end point of scientific discovery. It is therefore critically important to disseminate the findings in genetic association analysis to the downstream analysis and experimental work. From this perspective, Bayesian approaches are generally advantageous mainly because of their use of probabilistic quantification to summarize association results comprehensively. This point has been illustrated by the co-localization analysis of molecular QTL and GWAS signals, where most existing approaches (Giambartolomei et al., 2014, Hormozdiari et al., 2016, Wen et al., 2017) all require posterior probabilities from both complex and molecular trait-association analyses. For integrative analysis requiring results from genetic association analysis, e.g., SNP-level eQTL annotations, the probabilistic quantification of association results is more appropriate than the binary classification based on some stringent type I error threshold. This is because the latter approach fundamentally ignores the potential type II errors (which also contribute to the misclassifications) and can introduce severe bias in the integrative analysis.

It is worth pointing out that the equivalency of analysis results by individual-level and using summary-level data is based on the assumption that the correlation matrix between candidate variants, ***R***, is estimated accurately. The deviation from this assumption can cause noticeable discrepancy between the two types of analyses. We acknowledge that the estimation of ***R*** from an appropriate population is an important problem and refer the readers to some important recent works on this topic (Zhu et al., 2017).

## Web Resources

DAP-G software and tutorial, http://github.com/xqwen/dap/

GTEx data, http://www.gtexportal.org/home/datasets

Simulation data and code, https://github.com/xqwen/dap/tree/master/dap-g_paper/simulation Multi-SNP fine-mapping results of the GTEx whole blood data, https://github.com/xqwen/dap/tree/master/dap-g_paper/gtex_v6p

## Appendices

## Appendix A Details on Bayesian hierarchical linear model

Recall that we assume the linear model (1),

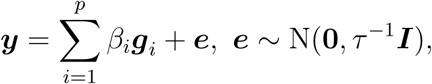

and define *γ_i_* = **1**(*β_i_*≠0). The *γ_i_*’s are assumed independent *a priori* with the following prior distribution,

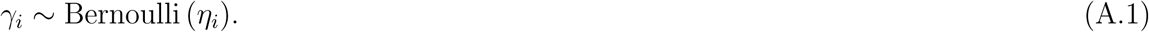

In case that an *m*-dimensional annotation, ***d***_*i*_, is available for each SNP *i*, we incorporate this quantitative information into the prior specification through a logistic function, i.e.,

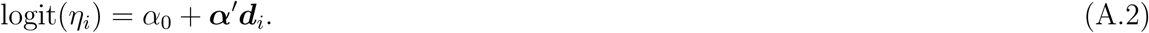

We estimate the enrichment parameters (*α*_0_, ***α***) from the observed data using an EM algorithm detailed in Wen (2016). In the absence of the annotation data, the logistic prior reduces to a single intercept term. The prior for the effect size parameter *β_i_* is assumed the following form

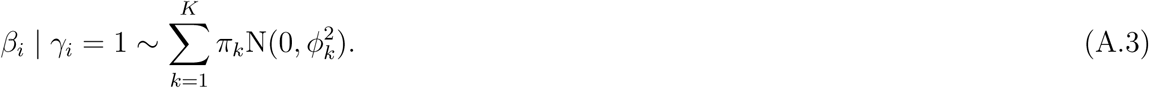

By including a grid of values for *ϕ_k_*, this mixture prior attempts to capture a spectrum of genetic effect sizes ranging from modest to strong. By default, *π_k_* is set to 1*/K*. It is also possible to estimate individual *π_k_* values by an EM algorithm (e.g., the one implemented in TORUS). The marginal priors on *β_i_*’s are known as spike-and-slab in the statistical literature.

Finally, we assume a Γ prior for the parameter *τ* that controls residual error variance in the linear model, i.e.,

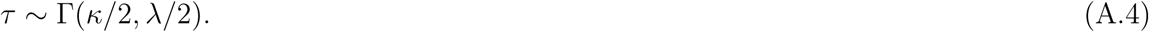

For inference, we assume the limiting form of this prior as *κ, λ →* 0.

## Appendix B Inference using summary statistics

### B.1 Sufficient statistics for likelihood computation

Our result is derived from the analytic expression of Bayes factors and their approximations in a general complex linear model system reported in Wen (2014), where the multiple linear regression model discussed in this paper is a trivial special case. Assuming for a given ***γ*** value, the linear regression model (1) is reduced to *q* assumed associated SNPs (i.e., *q* entries of the ***γ*** vector are 1) and we adjust ***G*** to denote the *q × q* design matrix specific to the value of ***γ***. Let the *q*-vector ***β*** denote the genetic effect sizes of the *q* SNPs. We assume a general prior distribution for ***β***,

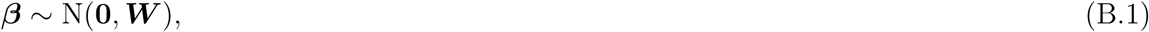

where ***W*** is a *q × q* positive semi-definite matrix. In this case, the Bayes factor can be computed by

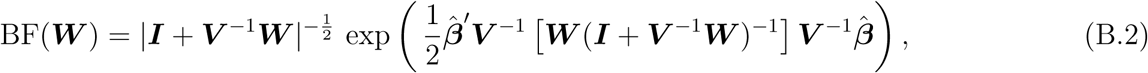

For multiple linear regression model, we note that

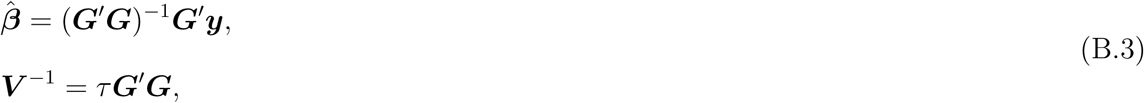

and (B.2) can be simplified to

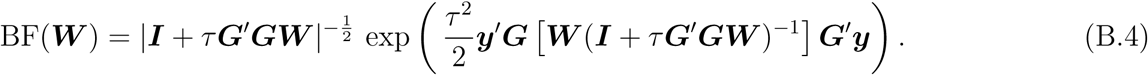

If *τ* is known, the above expression is exact, and the computation relies on the observed data only through the summary statistics (***G****′****y***, ***G****′****G***).

When *τ* is unknown, Wen (2014) shows the above analytic form becomes an approximation via Laplace’s method, i.e.,

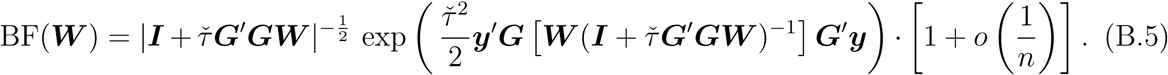

In particular, the point estimate *τ̆* is an affine combination of the MLEs of *τ* estimated under the null model (denoted by *τ̃*) and the full model (denoted by *τ̂*). More specifically,

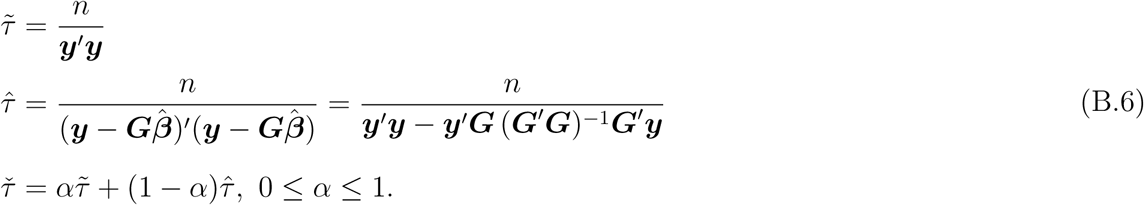

In other words, plugging in any value between *τ̂* and *τ̃* for *τ* corresponds to a valid Laplace approximation of BF(***W***). Note that this result is essential for the justification of the use of single-SNP testing *z*-scores when *τ* is unknown.

In addition to ***G****′****y*** and ***G****′****G***, estimating *τ̃* and/or *τ̂* requires two more summary statistics, sample size *n* and SST = ***y****′****y***, when *τ* is unknown. Thus, we conclude that the sufficient statistics required to compute the Bayes factors are (***G****′****y***, ***G****′****G****, n*, SST).

### B.2 Recovering sufficient statistics

When *τ* is known, single SNP testing *z*-statistics along with ***R*** and **Λ** are sufficient to recover the required sufficient statistics for Bayes factor computation, i.e.,

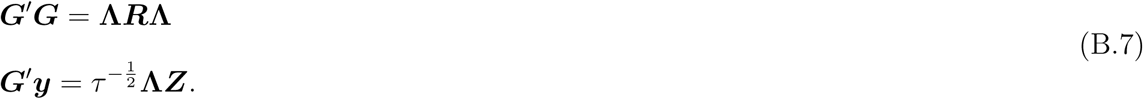

Here we focus our discussion on recovering the sufficient statistics in a realistic setting where *τ* is not known. In particular, we assume that for each SNP *i*, the effect size estimate

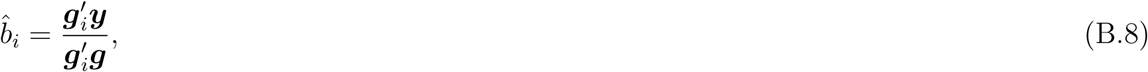

and its standard error, *ŝ_i_* = se(*b̂_i_*). Additionally, we only assume the knowledge of ***R*** (but not **Λ**), *n* and SST.

We show the following procedure can recover required sufficient statistics assuming ***R*** is accurate. For each SNP *i*,

1. Compute *z_i_* = *b̂_i_/ŝ_i_*
2. Compute *R*^2^ for the corresponding simple linear regression model by

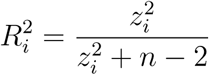
3. Find the estimated residual error variance from the corresponding simple linear regression model by

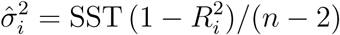
4. Compute 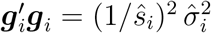
5. Compute *g*′*_i_y=b̂ _i_*∙*g*′*_i_g*

Subsequently, we obtain that

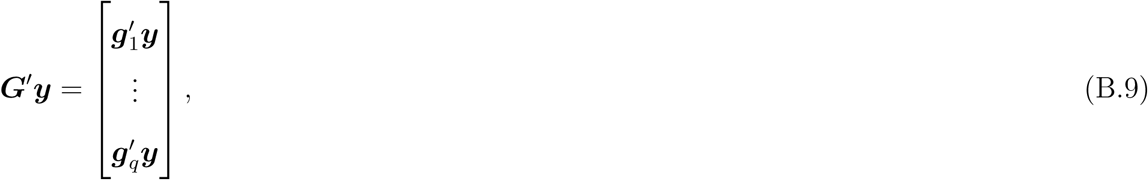

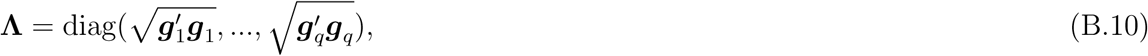

and

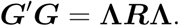

Consequently, any appropriate form of *τ* estimate can be obtained.

### B.3 Connection to previous results

In this section, we show that our results are connected to the existing literature, assuming *τ* is known.

#### B.3.1 Result for known *τ*

Assume that ***W*** is full-rank, it follows that

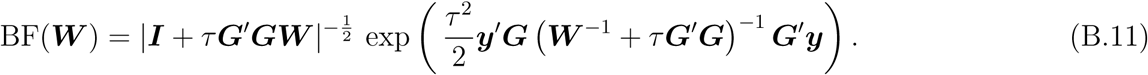

Plugging in Equation (B.7) results in

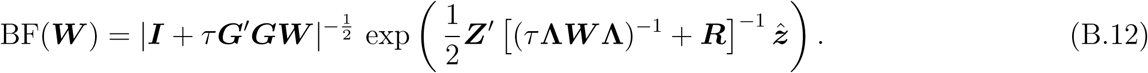

In particular, Chen et al. (2015) uses a specific form of prior, which scales the effect size of each SNP by its genotype variance and *τ*, namely,

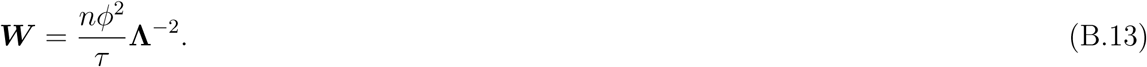

It follows from the Sylvester’s determinant theorem that

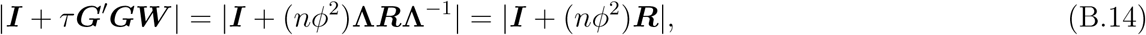

and

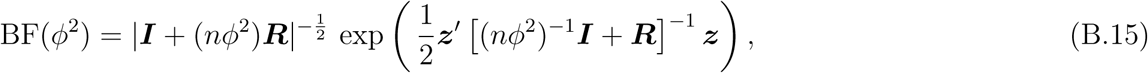

which only requires (***R***, ***z***) and is exactly the same result presented by Chen et al. (2015).

#### B.3.2 Result for unknown *τ*

Here we justify the use of the analytic form of Equation (B.15) when *τ* is not known. With the specific prior (B.13), the Bayes factor for a given *τ* value is

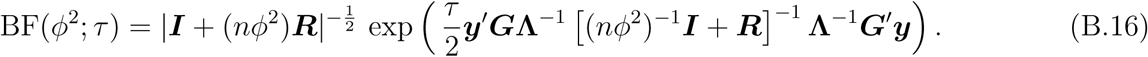

Wen (2014) shows that the desired Bayes factor with respect to arbitrary prior density *p*(*τ*) can be approximated by the Laplace’s method. The resulting approximation is given by

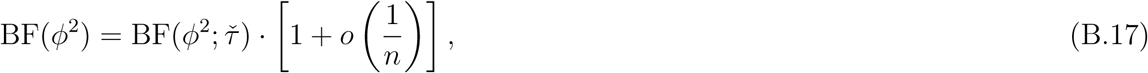

where *τ̆* can be any affine combination of *τ̃* (the MLE of *τ* from the null model) and *τ̂* (the MLE of *τ* from the full model). Note that the quadratic form,

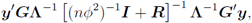

is positive definite, the approximate Bayes factor (ABF) is monotonically increasing with respect to the value of *τ̆*. More specifically, all valid ABFs justified by this approximation satisfy

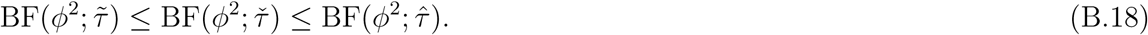

Alternatively, we can represent the ABF result as a function of multi-dimensional *z*-scores. First, we define a *p*-vector ***τ***:= (*τ*_1_*, τ*_2_*,…,τ_p_*), and denote

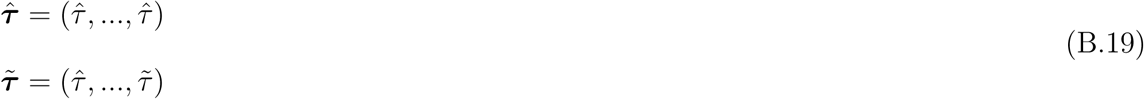

Let 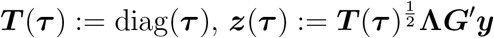, and we define

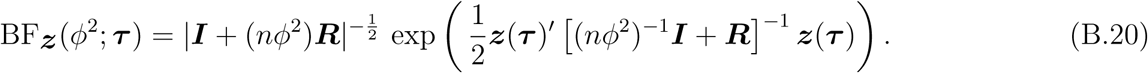

Here we attempt to link the analytic expression, BF***z***(*ϕ*^2^; ***τ****̆*), to the well-defined approximate Bayes factor BF(*ϕ*^2^; *τ̆*).

Let *f* (***z***) = ***z****′* [(*nϕ*^2^)^*−*1^***I*** + ***R***]^*−*1^ ***z*** = ***z****′****Az***, it follows that

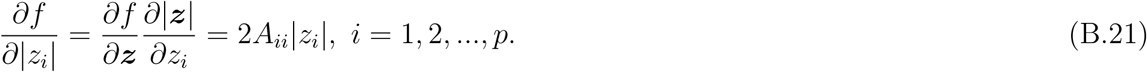

Because matrix ***A*** is positive definite, it follows that *A_ii_* = ***e****′_i_****Ae****_i_* > 0, ∀*i*, where ***e***_*i*_denotes the unit vector with the *i*-th entry being set to 1. Equation (B.21) indicates that *f* (***z***) is monotonically increasing with respect to each individual *|z_i_|*.

In practice, 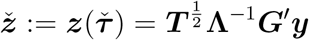 is used to evaluate BF***z***(*ϕ*^2^; ***τ****̆*), where ***T*** = diag(*τ̆*_1_*,…,τ̆_p_*) and each *τ̆_i_* represents the MLE of *τ* estimated from the simple regression model testing the association of SNP *i*. It should be clear that

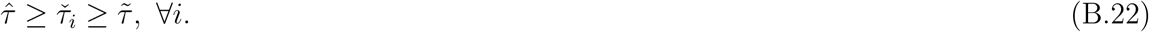

Let ***z****̂*:= ***z***(***τ****̂*) and ***z****̃*:= ***z***(***τ****̃*), it follows that

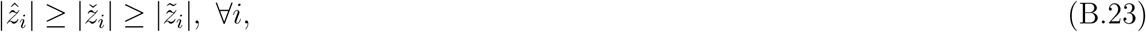

Consequently, it implies that

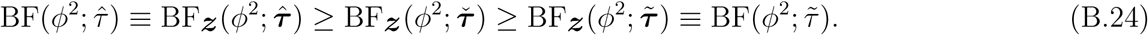

By the intermediate value theorem, there must exists 0 *≤ α ≤* 1, and

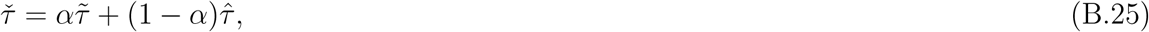

such that

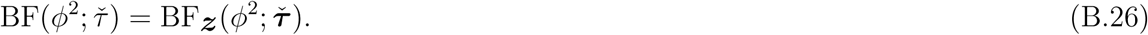

Therefore, BF***z***(*ϕ*^2^; ***τ****̆*) is valid approximation of Bayes factor under the prior (B.13) by the argument of Laplace’s method.

##### Numerical Illustration

If the association model under consideration (i.e., ***γ***) contains no true association signal, or the genetic effects of the suspected associations are small, we expect that *τ̂ ≈ τ̆ ≈ τ̃*. As a result, we also expect

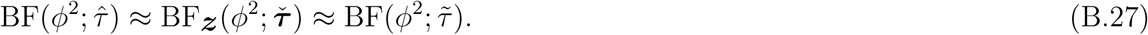

To illustrate, we simulate a quantitative traits for 343 individuals with 3 independent genetic variants, i.e.,

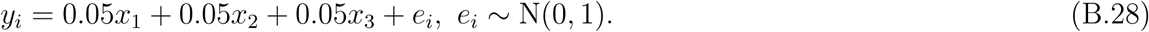

Assuming *ϕ*^2^ = 1, the comparison of the three approximate Bayes factors is shown in Table 2. In an alternative scenario where the data contain multiple modest to strong association signals, directly applying the *z*-score approximation can result in an equivalent *τ̆* value that severely over-estimates residual error, hence under-estimate the Bayes factor. To illustrate, we use the same simulated genotype data and the following linear model,

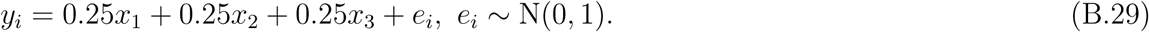

**Table 2:** Comparison of different approximate Bayes factors under weak association.

| $\log_{10} \text{BF}(\phi^2 = 1; \hat{\tau})$ | $\log_{10} \text{BF}_{\mathbf{z}}(\phi^2 = 1; \check{\tau})$ | $\log_{10} \text{BF}(\phi^2 = 1; \tilde{\tau})$ |
| --- | --- | --- |
| -0.776 | -0.822 | -0.870 |

As expected, the resulting approximate Bayes factors shows difference in order of magnitude (Table 3), albeit all approximations show overwhelming evidence of association.

**Table 3:** Comparison of different approximate Bayes factors under modest association.

| $\log_{10} \text{BF}(\phi^2 = 1; \hat{\tau})$ | $\log_{10} \text{BF}_{\mathbf{z}}(\phi^2 = 1; \check{\tau})$ | $\log_{10} \text{BF}(\phi^2 = 1; \tilde{\tau})$ |
| --- | --- | --- |
| 16.244 | 13.247 | 12.09 |

---

**Figure S1:**
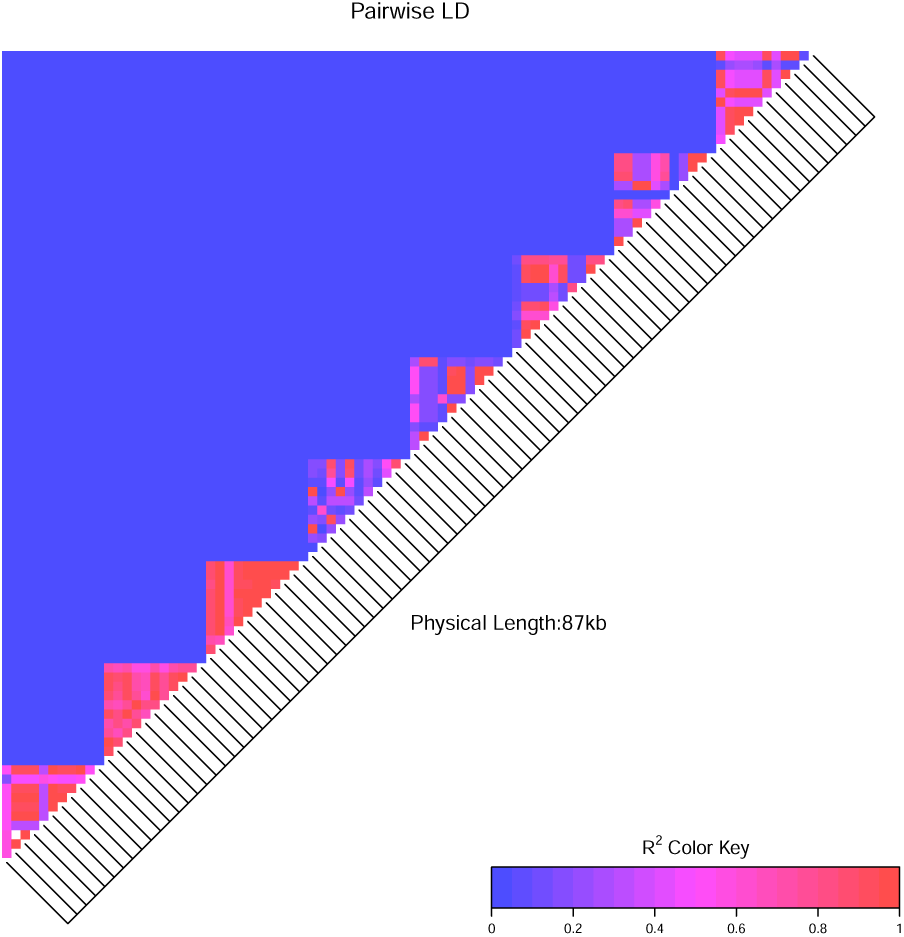
LD structures from 8 randomly selected blocks in the simulation study. *R*^2^ values are plotted for 88 SNPs from 8 artificially constructed blocks. The 8 blocks are randomly selected from a total of 91 blocks used in the simulations. All genotype data are real and from GUEVADIS study. By our construction, LD patterns within each block vary but the LD between blocks is consistently weak.

**Figure S2:**
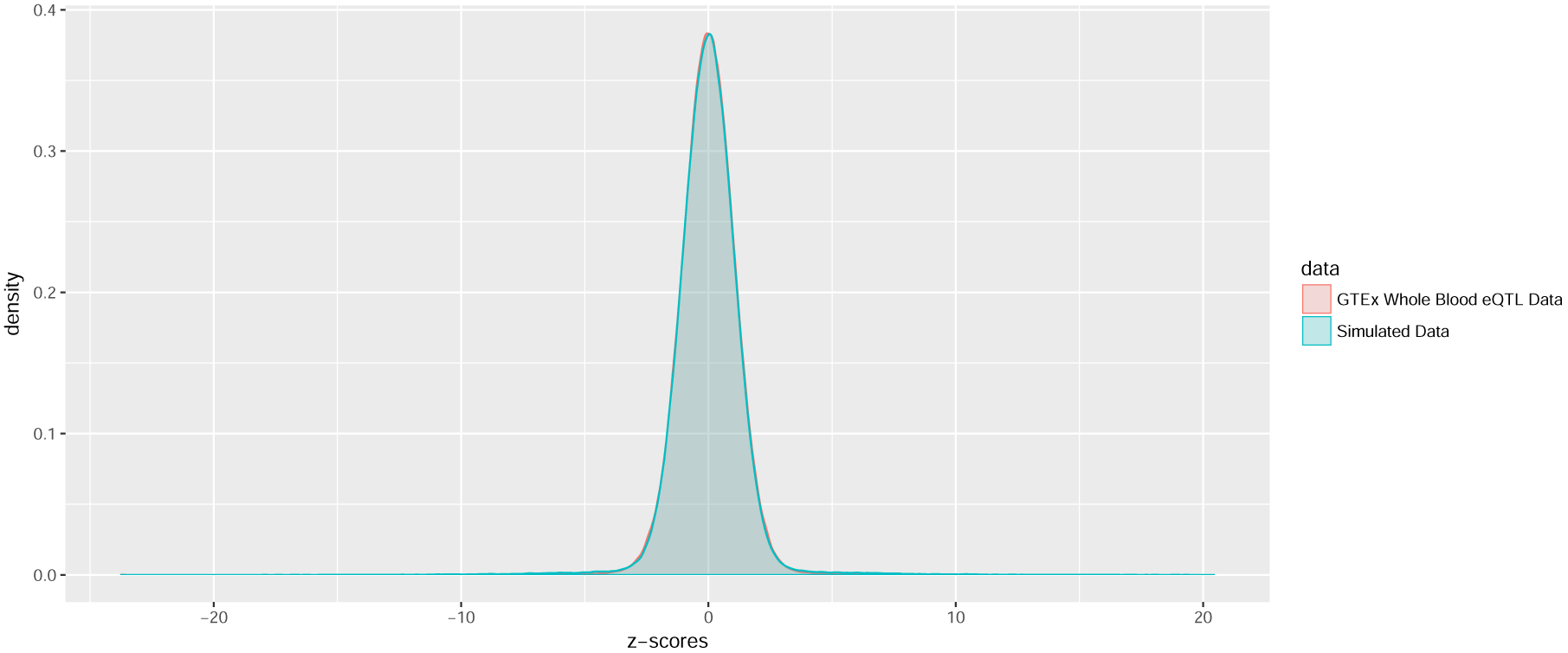
Comparison of single SNP *z*-scores between simulated data and GTEx whole blood eQTL data. The effect size parameters in the simulation studies are chosen to mimic the observed *cis*-eQTL data. The density of *z*-scores computed from the simulated data overlay almost entirely with the observed *z*-score distribution from the GTEx whole blood data.

